# *PHOX2B* polyalanine repeat mutation has a profound impact on the transcriptome of neuronal progenitor cells in Haddad syndrome

**DOI:** 10.1101/2025.10.15.682708

**Authors:** Tsering Stobdan, Vaishnavi Ventrapragada, Hang Yao, Dan Zhou, Ila Dwivedi, Daniel Lesser, Gabriel Haddad

## Abstract

Mutation in *paired-like homeobox 2B* (*PHOX2B*) is used as the diagnostic marker of Haddad syndrome (HS). The mutant gene/protein afflict neural crest cells during embryonic development which leads to congenital central hypoventilation syndrome (CCHS) and Hirschsprung’s disease (HSCR). Previous studies on HS and CCHS have mainly focused on the conformational dynamics of the mutant protein and have remained controversial. Here we performed RNA-sequencing on the patient derived neuroepithelial stem cells (NESCs), pertinent to the neurodevelopmental phenotype in HS, and found that the *PHOX2B-PARM* has a profound impact on the transcriptional profile of the cells. The single copy of *PHOX2B-PARM* in heterozygote cells were leading to >10 fold differentially expressed genes. In the patient cells there was a significant enrichment of genes related to *neuronal development* and *synapse organization* mainly driven by *L1CAM interactions* and *synaptogenesis signaling pathway*. Our result not only highlight the use of a suitable model of HS but also provide a clear path for future experimental validation and downstream targets with potential therapeutic values.

## Introduction

Haddad syndrome (HS) is a rare genetic disorder where individuals have both Congenital Central Hypoventilation Syndrome (CCHS), a severe genetic disorder of neurodevelopment resulting in disordered autonomic nervous system function, and Hirschsprung’s disease (HSCR), characterized by a lack of enteric nervous system ganglion cells, leading to bowel blockage^1^. Both CCHS and HSCR have no effective cure and have only surgical or symptomatic supportive therapies, such as positive pressure ventilation and/or phrenic nerve stimulation and surgical resection of the aganglionic segment, respectively^2^. Limited data from brain structure and function have revealed a wide range of observations that includes a normal brain^3,4^, non-specific, i.e., unrelated to autonomic nervous system dysfunction^5^ and a few with gray matter volume reduction^4,6^. Some of the related tissues include brainstem^7,8^, solitary tract^9-11^, locus coeruleus and arcuate nucleus^12-14^. Since RTN/pFRG, the brain region known to control breathing in mice^15^, is not well defined in humans, data obtained from mice models are not necessarily relevant.

The *paired-like homeobox 2B* (*PHOX2B*), a transcription factor involved in the development of the autonomic nervous system (ANS), was discovered as the defective gene in HS^16,17^ and the poly-Ala repeat mutation (PARM) and non-PARM (NPARM) are used as the diagnostic marker for HS and CCHS^16,18^. Note that the wildtype *PHOX2B* harbors 20 consecutive alanine residues, and in PARM an addition of 4-13 alanine residues is observed in the disease, with the longer expansions correlating with a more severe clinical phenotype^19-21^. The molecular dynamics of the mutant PHOX2B protein, i.e., aggregate formation and cytoplasmic mis-localization^22-24^, has long been the subject of extensive investigations, but remained controversial^25^.

At the molecular, biochemical and structural levels, HS has two critical outstanding and pressing issues^26^: a) how *PHOX2B-PARM* or general poly-A expansion functions, and b) how mutation in *PHOX2B* gene leads to HS, i.e., what are the downstream targets of *PHOX2B-PARM* that lead to HS? There has been continuous progress made in understanding the structural dynamics^25^ and the molecular mechanisms underlying *PHOX2B-PARM*. For example, *PHOX2B-PARM* is known to function as a toxic gain of function, expressed in an autosomal dominant manner. However, investigations to identify the downstream targets of *PHOX2B-PARM* that can directly impact HS, remain scarce.

Previous studies using mice or in-vitro cell models have helped reveal critical information^11,15^. In mice, RTN is identified to play a critical role in perinatal respiratory control and its development was reportedly prevented in *Phox2B*^*27Ala/+*^ mice^15^. However, observing a hypoplastic RTN in CCHS is implausible when its location or existence is also controversial^27,28^. In cell lines previously used, the endogenous expression of *PHOX2B* was unusual, i.e., either very high, as in the case of neuroblastoma^29^ or completely absent, as known in HeLa^23^. In a model that could mimic HS, it would be useful to have a similar genetic background, a similar spatiotemporal expression, or more importantly in the heterozygous state, as is the case in HS/CCHS patient. To address these issues, here we reprogram HS patients’ cells into iPSCs and differentiated them into NESCs. Furthermore, our RNA-seq analysis of the *PHOX2B* expressing NESCs have shown a profound impact of *PARM* on the transcriptome profile of early neuronal cells, pertinent to an early developmental stage in the human embryo. Due to the clinical importance of *PHOX2B-PARM*, our study has substantial medical relevance and will provide unique insights into early brain development.

## Materials and methods

### Patient samples

Individuals clinically suspected or diagnosed with CCHS were referred for genetic testing by the co-investigators at Rady Children’s Health, San Diego. A combination of next generation sequencing and sanger sequencing was used to cover the full coding regions of PHOX2B gene (https://www.preventiongenetics.com/). Based on the clinical findings there was 99% chance for detecting PHOX2B mutation in affected individuals. Once confirmed, we recruited the probands and their family members from Rady Children’s Hospital, San Diego and acquired their blood/cells (confirmed PHOX2B-PARM genotypes in probands). Informed consents were obtained from the parents of the affected children and the volunteering family members. Institutional review board approved the study.

### Reprogramming to iPSC

Blood sample was collected and subsequently peripheral blood mononuclear cells (PBMCs) were isolated from the blood. Patient derived PBMCs were reprogrammed to iPSCs by the CytoTune-iPS 2.0 reprogramming kit (ThermoFisher Scientific, Cat# A16517) according to the manufacturer’s manual. Culture plates were pre-coated with corning matrigel hESC-Qualified Matrix (1.25%) and kept in incubator containing 5% CO2 at 37°C. The transduced cells, i.e., iPSCs, were cultured with mTeSR1 medium (STEMCELL Technologies).

### Neuroepithelial stem cell (NESC) Generation

The protocol to generate NESCs was previously mentioned^28,30^. Briefly, iPSCs were differentiated to NESC by neural lineage specifications induced by inhibition of transforming growth factor β (TGFβ, SB431542) and bone morphogenic protein (BMP, LDN), and by induction of WNT (CHIR99021) and Hedgehog signaling pathways (Purmorphamine)^31^.

### eGFP tagging of PHOX2B expressing cells

The two critical limitations to isolate cells expressing *PHOX2B* includes, a) *PHOX2B* itself being a transcription factor is localized inside the nucleus and it does not have a proxy surface antigen for antibody staining, and b) the *PHOX2B* expressing cells in the NESCs were <1%, and paraformaldehyde fixation and staining were inefficient.

Therefore, we tagged PHOX2B-expressing NESCs with eGFP. For this we synthesized 1723bp construct (GenScript) that constituted a) a 1000bp upstream sequence of *PHOX2B* start codon (Phox2b-Prom), b) kozak sequence for efficiency and accuracy of translation initiation and c) protein coding eGFP sequence (GenScript). The 1000bp upstream of PHOX2B start codon was previously identified to have the complete promoter region^32^. The complete construct i.e., Phox2b-Prom-eGFP, was then cloned into lentivirus pGenLenti using CloneEZ (GenScript Cat# L00339). The packaging, concentration, and titration of lentiviral particles was carried out at viral vector core facility at Sanford Burnham Prebys. We followed a standard protocol for lentivirus transduction into NESCs that uses polybrene to enhance the efficiency. Puromycin selection confirmed successful infection of NESCs. Cells expressing eGFP was observed under Sterling X1 Spinning Disk, a multifunctional microscope (Nikon Imaging Center at the UCSD). The eGFP expressing cells co-localizes with PHOX2B antibody (sc-376997, SC Biotech. Inc.) staining (Fig 1D).

**Figure 1:**
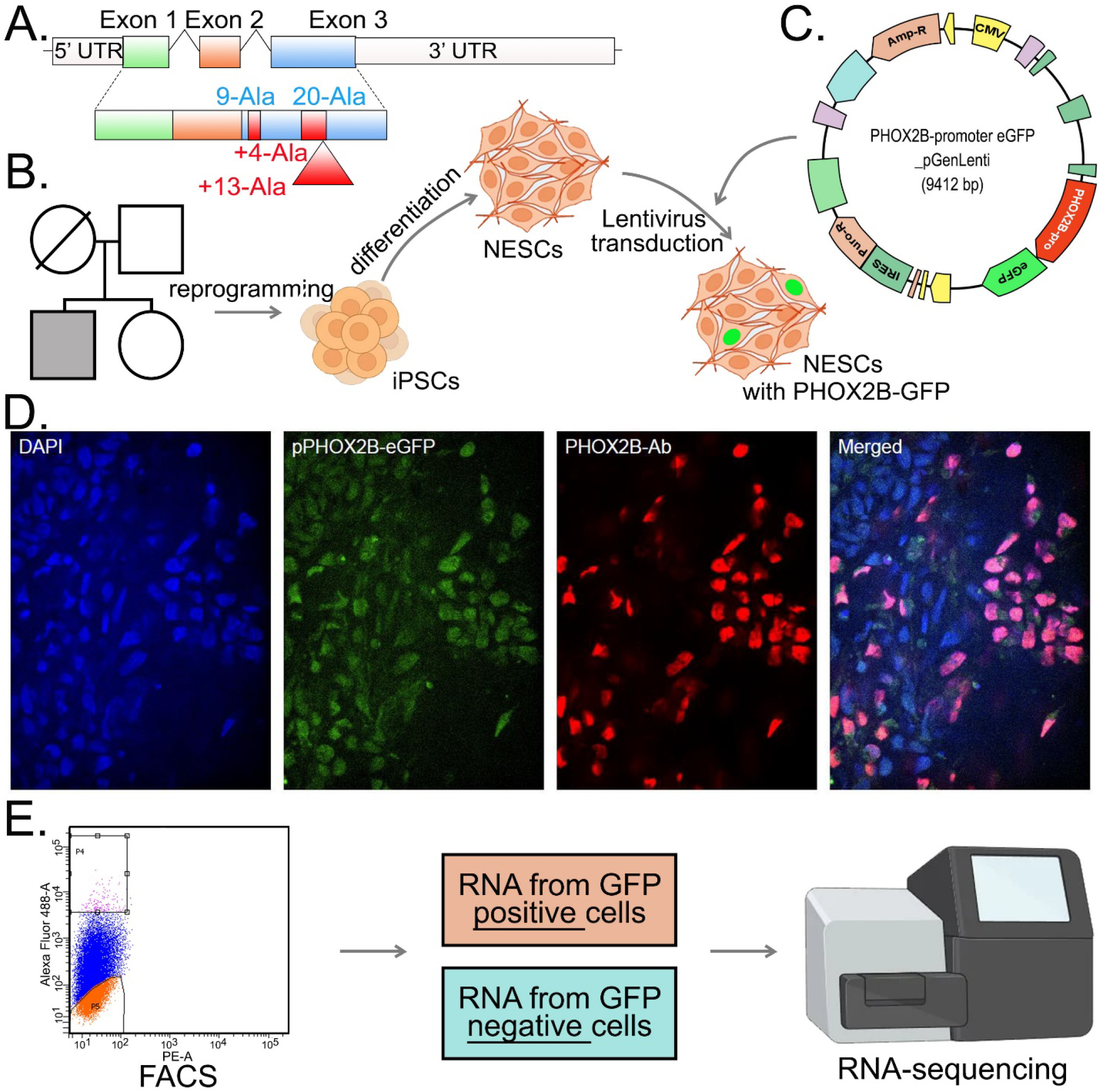
(A) The *PHOX2B* gene depicting PARM. (B) Generation of NEPCs from patient (Proband in gray box) and control. PBMCs were reprogrammed to iPSCs and differentiated to NEPCs. The cells were then infected with lentivirus carrying eGFP carrying *PHOX2B* promoter sequence. (C) *PHOX2B* promoter-eGFP (1723 bp) cloned in pGenLenti cloning vector (7689 bp) and selected for Ampicillin resistance. (D) Cells expressing eGFP derived by *PHOX2B* promoter activity co-localize with cells stained with PHOX2B antibody. (E) PHOX2B^+^ and PHOX2B^-^ cells were separated using FACS followed by RNA-sequencing.

### Fluorescence-activated cell sorting (FACS) to separate PHOX2B expressing cells

Next, we separated PHOX2B expressing cells using FACS (Fig 1E). Briefly, after removing the old media, cells were treated with accutase (50% with PBS) for 15mins at 37°C (intermittently the plates were gently swirled). Dissociated cells were collected in 15ml tubes and washed 1x with PBS, centrifuge for 3 mins at 2500rpm and resuspended in ice cold sort buffer (PBS with 1%FBA) at 5×10^6 cells per mL. Cells were filtered through 35µm nylon cell strainer (Falcon# 352235) and was kept on ice to proceed for FACS. Cells were analyzed using FACSDiva software (BD Biosciences) and sorted with FACSInflux, FACSAria Fusion or FACSAria2 (BD Biosciences).

We collected four groups of cell fractions i.e., GFP ‘+’ fraction from probands (P2BM) and controls (P2BW), and GFP ‘-’ from probands (NP2BM) and controls (NP2BW). The GFP ‘+ve’ cells were the top 1-2% of the clusters and GFP’-ve’ cells constituted cluster of cells that overlay the negative control i.e., cells that were not transduced with Phox2b-Prom-eGFP.

### RNA isolation and high throughput RNA sequencing (RNA-seq)

After FACS the cells were centrifuged for 3 mins at 2500rpm. The supernatant was removed leaving only 100ul of the sort buffer. We used Direct-zol RNA Microprep Kits (Zymo Research Cat#R2051) for RNA isolation. RNA samples were submitted to Institute for Genomic Medicine (IGM), UCSD, for quantity and quality check, library preparation and sequencing. Agilent Tapestation instrument was used for quantity and quality assessment. Samples with RNA Integrity Numbers (RIN) score□>□7 were used. TruSeq mRNA Stranded Library Prep Kit (Illumina) was used for library preparation. Sequencing was performed on the Illumina NovaSeq 6000 to produce 100 base pair, paired end reading.

### Differential gene expression analysis

The raw FASTQ file obtained from IGM was uploaded to Illumina BaseSpace Sequence Hub using the command line interface. RNA-Seq Alignment workflow version 2.0.2 was used to perform mapping using the STAR aligner^33^, quantification of reference genes and transcripts using salmon^34^. Homo sapiens UCSC hg19 was used for reference gene alignment. The Illumina® DRAGEN RNA-Seq differential expression application version 1.0.1 was used to identify differentially expressed genes (DEGs) which does differential expression analysis of reference genes with DESeq2. We used DEGs with adjusted p-value <0.05 for downstream analysis.

### Gene ontology and gene set enrichment analysis

The gene ontology, gene list enrichment analysis and candidate gene prioritization were done using online tools at www.geneontology.org and https://pantherdb.org/. This was further validated using ToppGene Suite^35^. Ingenuity Pathway Analysis (IPA; Qiagen Inc.,) was used to identify molecular function from the top DEGs. An assigned activation z-score from IPA reveals if the pathway is activated or inhibited.

### Boolean implications analysis

We utilized publicly available GTEx Analysis V8, that constituted n=17382 human RNA-seq dataset, to perform Boolean implication analysis^36^. We used *PHOX2B* as the seed gene. The known targets of *PHOX2B* were obtained from the literature.

## Results

### Reprogramming, differentiation and separation of HS PHOX2B^+^ cells

After obtaining approval from the institutional review board (see Methods), we recruited probands, with confirmed PHOX2B-PARM, and their family members from Rady Children’s Health, San Diego. The peripheral blood mononuclear cells (PBMCs, CD34^+^ cells) from probands and controls were successfully reprogrammed to iPSCs, which were then differentiated into neuroepithelial stem cells (NESCs, Fig. 1B). PHOX2B antibody (sc-376997, SC Biotech. Inc.) staining confirms that <1% of cells express PHOX2B. In order to isolate RNA from this rare cell population, we tagged PHOX2B-expressing NESCs with eGFP by transfecting pGenLenti that constituted *PHOX2B* promoter-eGFP (Fig 1C, details in Methods Section). Co-localization of GFP with PHOX2B antibody staining confirms PHOX2B’+ve’ cells. Next, we separated PHOX2B expressing cells using FACS (Fig 1E). We were able to collect four groups of cell fractions for RNA sequencing, i.e., GFP’+ve’ fraction from probands (P2BM) and controls (P2BW), and GFP’-ve’ from probands (NP2BM) and controls (NP2BW).

### Novel downstream targets of PHOX2B

In order to confirm that the GFP signal represents the expression of *PHOX2B*, we compared RNA-seq data of *PHOX2B’+ve’* cells and *PHOX2B’-ve’* cells of the control group. Interestingly, out of the known *PHOX2B* target genes, i.e., *PHOX2A, RET, TH, DBH* and *ASCL1*, only *PHOX2A* differed significantly between PHOX2B’+ve’ and PHOX2B’-ve’ (Fig. 2A and 2B). Although, *RET* and *TH* showed a positive relationship, no change was noted for *DBH* and *ASCL1* (Fig. 2B). Similarly, some of the known targets, e.g., *TLX2*^37^, were not even expressed at this stage. Furthermore, our data also reveal 228 differentially expressed genes (DEGs) between *PHOX2B’+ve’* cells and *PHOX2B’-ve’* cells (adjusted P<0.05, Fig 2A), indicating that they have a potential relation with *PHOX2B*.

**Figure 2:**
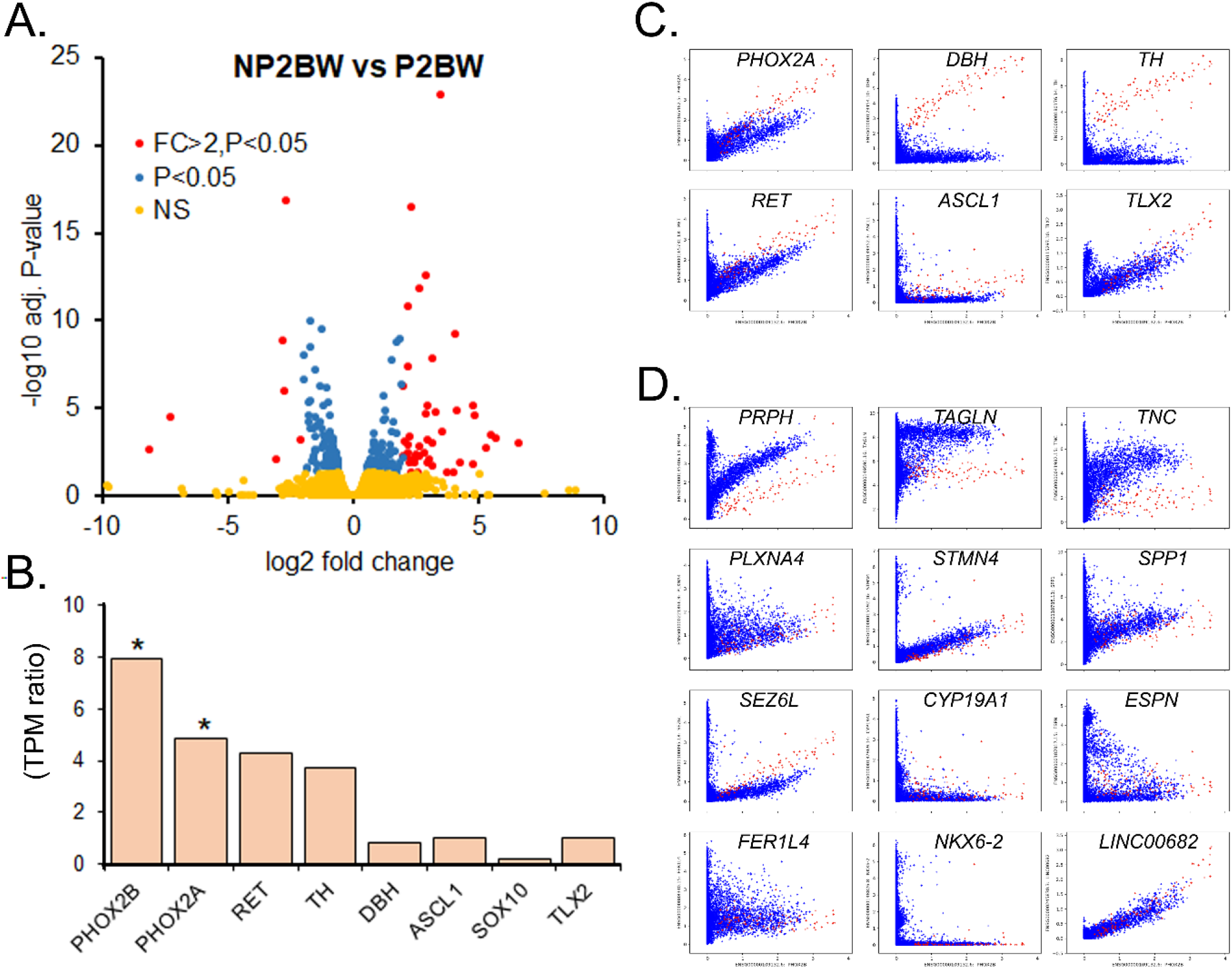
Novel downstream targets of PHOX2B. (A) Volcano plot depicting DEGs due to normal *PHOX2B* expression. (B) Ratio of transcript per million (TPM) of the known PHOX2B target genes, between PHOX2B’+ve’ i.e., P2BW and PHOX2B’-ve’ i.e., NP2BW samples. (C) Boolean analysis of the known *PHOX2B* target genes. The x-axis is the TPM for *PHOX2B* and y-axis is the TPM values for respective genes. The red dots are the cells/tissues that has a relatively stronger correlation between *PHOX2B* and its know targets genes i.e., *DBH* and *TH*. (D) Boolean analysis of some of the DEGs in the normal PHOX2B’+ve’ compared to PHOX2B’-ve’.

In order to validate these findings, we performed a Boolean analysis between *PHOX2B* and the known target genes, and with the top 53 DEGs from the current study, i.e., genes with >2 log_2_ fold change and adjusted P<0.05. As anticipated, we see a direct correlation between *PHOX2B* and *PHOX2A* (Fig 2C). We also noted a similar correlation for *PHOX2B* with *RET* and *TLX2* (Fig 2C). However, a direct correlation of *PHOX2B* with *DBH* and *TH* were limited to a minor cell population (Fig 2C). Interestingly, of the 53 genes we tested, >50% showed a correlation that was in par with the known *PHOX2A, RET* and *TLX2* (Fig 2D), clearly depicting a direct relationship of these genes with *PHOX2B*. Additionally, we also performed Boolean implications for genes that have >3 log_2_ fold change but did not reach the P-value cutoff, due to low TPM count or higher variability. Remarkably, a long non-coding RNA, *LINC00682*, showed an ideal Boolean relationship with *PHOX2B*, better than any of the known targets such as *PHOX2A* (Fig 2D). Our data identify novel target genes that merit further validation.

### *PHOX2B-PARM* induces robust transcriptional change

We report here a profound impact of heterozygote PHOX2B-PARM on the transcriptome profile of NESCs. The overall analysis of the RNA-seq using principal component analysis (PCA) reveals a batch effect where PC1, at 38.4% of its variance, cluster samples based on the two batches of RNA sequencing (Fig. 3A). Interestingly, PC2 (at 24%) clearly separates PHOX2B expressing patients cells from rest of the samples, including from the patients’ cells that do not express PHOX2B (Fig. 3A). It was the PC2 and PC3 that separate samples based on the PHOX2B protein expression where PHOX2B’+ve’ are grouped separately from PHOX2B’-ve’ (Suppl Fig. 1).

**Figure 3:**
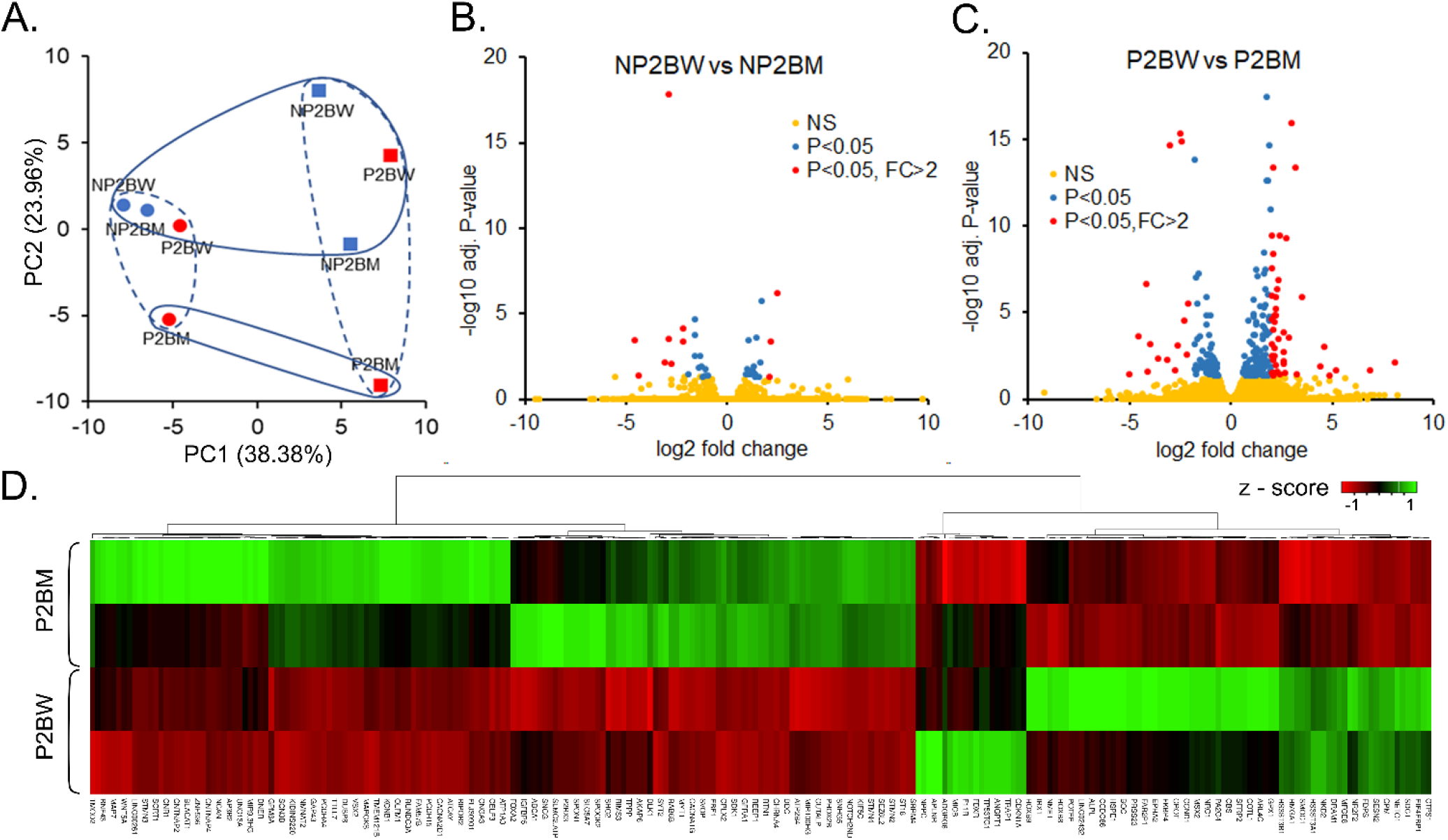
*PHOX2B-PARM* induces robust transcriptional change. (A) PCA depicting clustering of *PHOX2B-PARM*. (B) DEGs between NP2BM vs NP2BW. (C) DEGs between P2BM vs P2BW. (D) Heatmap depicting DEGs between P2BM vs P2BW.

A direct comparison between controls (P2BW) and patients (P2BM) reveals 242 DEGs. This number was >10 times the number of DEGs (n=23) detected for a baseline comparison, i.e., PHOX2B’-ve’ controls (NP2BW) vs PHOX2B’-ve’ patients (NP2BM) (adjusted P<0.05, Fig 3B and 3C). Interestingly, n=13 DEGs were common in both comparisons.

We also looked at the fold change of the DEGs, between P2BW and P2BM. There were 59 DEGs that had >2 log_2_ fold change and 17 DEGs with >3 log_2_ fold change (Fig 3C). In contrast, between NP2BW vs NP2BM there were only 11 genes that had >2 log_2_ fold change and only two genes that had >3 log_2_ fold change (Fig 3B). There were also 5 genes that were common in both comparisons. Taken together, the single copy of *PARM* induces a significant impact on the transcriptome profile of *PHOX2B* expressing NESCs.

### *PHOX2B-PARM* affect genes involved in neuronal development

We used RNA-seq data of NEPC stage to address the hypothesis that the presence or expression of *PHOX2B-PARM* during early neuronal development dysregulates the transcriptome profile which in turn affects the cellular composition leading on to spatial and temporal changes in HS patients during early brain development. The Fold-Change-Specific Enrichment Analysis (FSEA) of the DEGs between patients and controls PHOX2B’+ve’ cells was performed using the complete gene ontology (GO) annotation datasets. Remarkably, all the top GO terms for biological processes (GO-BP) were related to neuronal development and synapse organization (Fig 4A). For example, some of the GO-BP terms for *neuronal development* were generation of neurons, neuron differentiation, neuron development, and for *synapse organization* were synaptic signaling, anterograde trans-synaptic signaling, chemical synaptic transmission (Fig. 4A). Interestingly, among the constitutive genes that represent these GO terms, i.e., 64 genes related to generation of neurons and 29 genes related to synapse organization, >75% i.e., 49 and 25, respectively, were upregulated in the patients. The top cellular components (GO-CC) impacted by these DEGs were axon, neuron projection, synapse, and presynaptic membrane (Fig. 4B).

**Figure 4:**
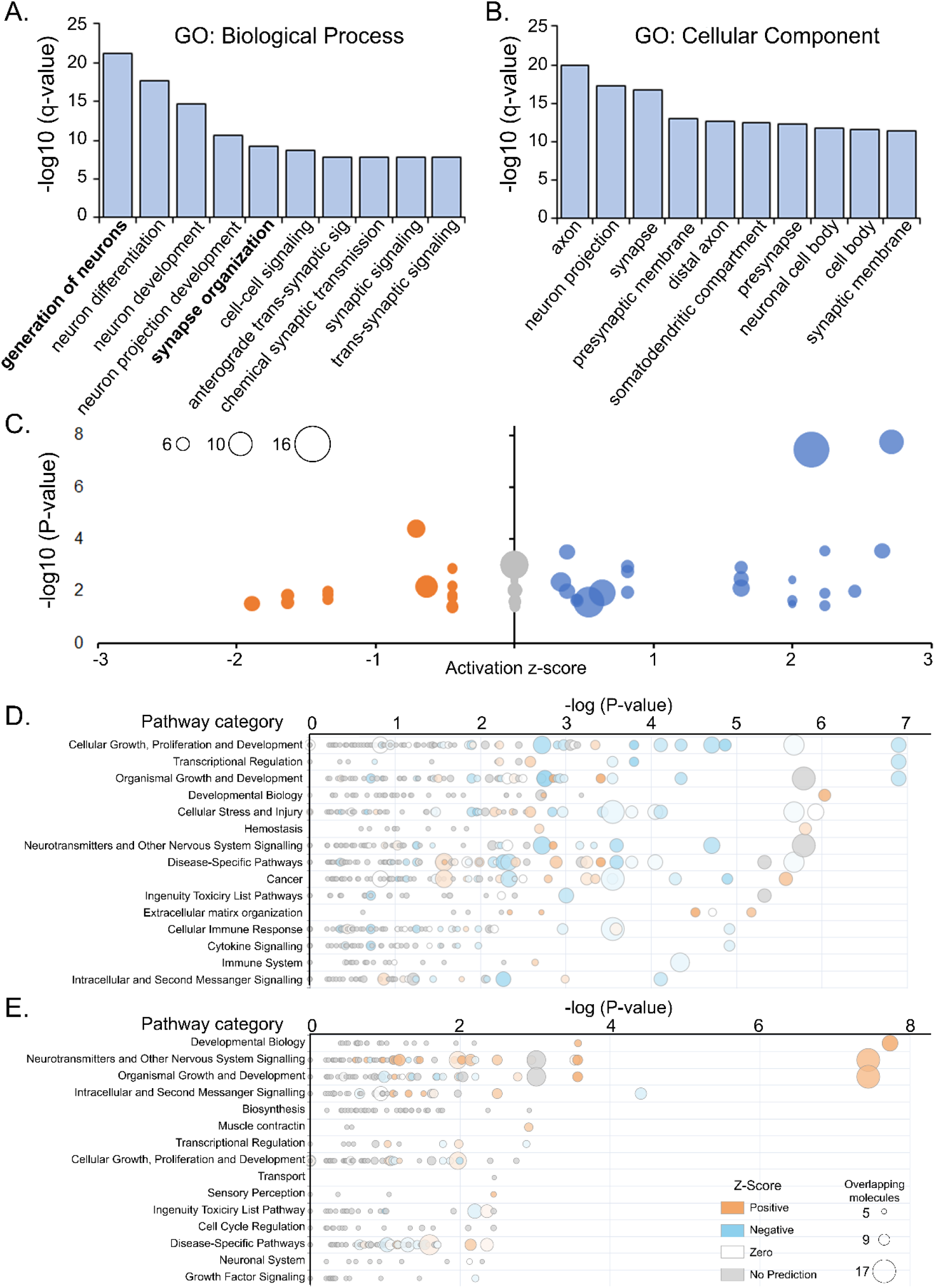
IPA of the DEGs. (A) GO-biological process and (B) GO-Cellular component of the top DEGs between P2BM vs P2BW. (C) Volcano plot depicting the activation score for different biological processes. A negative activation score indicated the biological process to be ‘inhibited’. (D) The biological pathway impacted in the NP2BM vs NP2BW. (E) The biological pathway impacted in the P2BM vs P2BW.

Our approach of using PHOX2B’+ve’ cells also provides us the opportunity to study the role of normal *PHOX2B*. Here we performed FSEA of DEGs from normal *PHOX2B* expression i.e., between PHOX2B’+ve’ and PHOX2B’-ve’ from controls. Although, once again some of the pertinent GO terms were related to generation of neurons, the candidate DEGs here were different (Suppl. Fig 2). In parallel some of the unique biological processes were tube development, positive regulation of developmental process, glutamatergic synapse and growth factor binding and DNA-binding transcription activator activity.

### *PHOX2B-PARM* has a major impact of L1CAM interactions and synaptogenesis signaling pathway

We used ingenuity pathway analysis (IPA) to identify signaling pathways impacted by *PHOX2B-PARM*. Some of the top impacted pathways were L1CAM interactions, synaptogenesis signaling pathway, adipogenesis pathway, NCAM signaling for neurite outgrowth and SNARE signaling pathway (Fig 4C). The activation z-score reveals that all these pathways, except for the adipogenesis pathway (z-score = -0.71), were activated (Fig 4C). For example, L1CAM interactions (activation z-score = 2.714) and synaptogenesis signaling pathway (z-score = 2.138) have most of the constitutive genes i.e., *L1CAM, NCAM1, NFASC, SCN1A, SCN3B* and *TUBB4A*, upregulated in the patients (P2BM).

The IPA predicts that the disease related biological functions activated by the DEGs were development of neural cells (activation z-score = 3.37) and the biological function inhibited were gastrointestinal tumors (activation z-score = -1.03).

We also utilized IPA to study the role of normal PHOX2B activation and among the top two impacted pathways the ‘transcriptional regulatory network in embryonic stem cells’ was inhibited (z-score = -1.508) and the ‘L1CAM interactions’ was activated (z-score = 2.333).

The overall biological process impacted by normal *PHOX2B* was related to inhibition of cellular growth, proliferation and development, transcriptional regulation and development (Fig 4D) and by*PHOX2B-PARM* was related to the activation of developmental processes, and neurotransmitters and nervous system signaling (Fig 4E).

## Discussion

Here we presented RNA-seq data of *PHOX2B*-expressing neuronal progenitor cells, comprising a copy of *PHOX2B-PARM*, recapitulating the transcriptome profile of the early neurodevelopmental stage in an HS patient. To our knowledge this is the first study on patient-derived *PHOX2B*-expressing neuronal progenitor cells. We identified downstream targets of *PHOX2B-PARM* and anticipate that some of these targets may have therapeutic potential to treat HS or CCHS.

The encouraging results from earlier studies in mice and human cell line were hindered for their translation potential due to limitations of a suitable model system. This includes, a) the *PHOX2B* expressing cell population being rare and transient, b) the spatiotemporal dynamics of cells during early neurodevelopment, c) fundamental limitations of murine models for human diseases and, d) the heterozygous state of *PHOX2B-PARM* involving a toxic gain of function mechanism. By using patient-derived NESCs combined with FACS, we are able to address all these limitations and obtained results from cells closely mimicking a critical stage of HS pathogenesis i.e., the early neuronal development in humans. The FACS separation of rare PHOX2B expressing cells is also confirmed by the overexpression of *PHOX2A*, the paralog gene directly known to associate with *PHOX2B* expression^38^. From our RNA-seq and Boolean analysis results we noted that the expression of some of the well-known target genes, e.g., *DBH*^39^, may not directly associate in all cell types. This is particularly surprising in light of the studies where *DBH* has been used as a proxy for *PHOX2B*^39^. Interestingly, at the same time Boolean analysis also reveals a better correlation for *PHOX2B* targets genes detected in cells related to HS i.e., *RET*^40^ and *TLX2*^37^. Therefore, our Boolean analysis has not only supported this notion but has also identified novel targets, that have a better correlation, and therefore offering a distinct advantage over previously used model systems.

Functionally, *PHOX2B-PARM* has been shown to dimerize with *PHOX2B*^24,25^ and because the mutation is outside the homeodomain region, we anticipate the protein retaining its DNA binding property. To this end, there have been significant attempts made to understand its structural dynamics. As a result there are numerous studies showing aggregate formation of *PHOX2B-PARM* ^21,22^, and there are disagreements as well, including ours (unpublished data), where we and others have failed to detect any aggregates^25,41^. Plausibly, unlike in previous studies where the mutant gene/protein was highly expressed e.g., in high *PHOX2B* expressing neuroblastoma^29^ or driven by expression vector^23^, the mutant protein when expressed at an endogenous level fails to aggregate. Indeed, in a defining aspect of its structural dynamics a recent study has uncovered *PHOX2B-PARM* triggering rapid phase transition into solid condensates that also captures the wild-type *PHOX2B*^25^. Due to the significant role of transcriptional condensates in transcription regulation^42^, we believe the pathogenic *PHOX2B-PARM* condensates will have a significant impact on the transcription profile.

In the current study we have shown a distinct transcriptome profile of the patient-derived PHOX2B-expressing NESCs as depicted by the patients’ samples from different batches clustering together and separate from other cell populations. Interestingly, there are only three previous studies, to date, that have done transcriptome profile to study HS/CCHS^29,41,43^. This is despite the transcriptome profiling being one of the most utilized approaches to investigate human diseases^44^. Additionally, only one of these studies used patient-derived cells^43^. The RNA-sequencing in that study was performed on a relatively mature neuronal cell population, contrary to the early neurodevelopmental stage impacted in HS/CCHS. Interestingly, similar to our current results, the major biological process impacted was ‘neuronal development’, even though the only shared gene between the two studies was *neuropilin and tolloid like 1* (*NETO1*) that was similarly downregulated in the HS/CCHS, in both studies. The expression of *NETO1* in the normal cells is upregulated during development and reported to have a role in hippocampal network growth^45^, there is little information about its role during brain development.

The other two transcriptome profile studies have used cells transfected with plasmid carrying mutant *PHOX2B-PARM* and have not taken into account the critical heterozygote state that leads to its toxic gain-of-function property^25,29,41^. However, the study that used a neuroblastoma cell line, the *teneurin transmembrane protein 4* (*TENM4*), involved in establishing proper neuronal connectivity during development, was common in our study and is upregulated is HS/CCHS^29^. In the other study, snRNA-seq was performed on the 60-day brainstem organoid of transfected control hPSC cell line^41^. Interestingly, regardless of the cell-type and approach, the biological process impacted in the mutant cells was related to ‘*neuron development*’ which we clearly noted in our current study. Furthermore, the organoid study also reported a significant impact on the cellular composition of the 60-day organoids^41^. We believe that our data complemented this study by providing a detailed insight of an earlier time point of brain development that impacts cellular composition at a later stage in development.

Recent studies have shown that the morphology, connectivity and function of neurons, i.e., their phenotypes, are dictated, to a large extent, by the transcriptome profile during an early stage, i.e., neuronal progenitor cells^46^. It is thus possible that the profound impact of *PHOX2B-PARM* on overall transcriptome holds key information in understanding HS pathogenesis. As the number of DEGs between control and patient was 10 times higher than the baseline comparisons, i.e., between non-PHOX2B expressing cells, it is likely that these DEGs contribute to remodeling of large portions of the neuronal cells over the course of development. Of relevance as well, the biological processes impacted were all related to generation of neurons and synapse organization, similar to previous studies^29,41,43^. Remarkably, multiple genes involved in these biological processes were upregulated in mutant cells. For example, the *L1CAMs* (L1 family of cell adhesion molecules)^47^, *NFASC* (Neurofascin)^48,49^ and *NCAN* (neurocan)^50^ have been implicated in processes integral to nervous system development, including neurite outgrowth, neurite fasciculation and inter neuronal adhesion, were all upregulated. This in turn ‘*activates*’ the existing biological processes, as depicted by IPA. These results led us to believe that the transcriptome change by *PHOX2B-PARM* disrupt the normal cell lineages in mutant cells, and consistent with this premise, such development, at least in part, could lead to premature ‘generation of neurons’ and ‘neuron differentiation’ of the NESCs, as indicated by some of the top GO terms.

Finally, for a disease like HS, where the causative gene is well established, we believe the shortest path to understand its molecular mechanism is the identification of the dysregulated downstream targets. By overcoming some of the critical limitations, we now have the RNA-seq data revealing *PHOX2B-PARM* targets molecules. Additionally, our current approach is downstream of current controversy related to the conformational changes^21,22,25^, and provides a clear path for future experimental validation. Since the cells that we analyzed are at the core of HS i.e., patient derived cells expressing the key gene (heterozygous *PHOX2B-PARM*), we believe our current study is a significant step towards understanding the functional consequences of *PHOX2B-PARM* in HS. Future studies on spatiotemporal development will be valuable in gaining a deeper understanding of HS pathogenesis.

## Acknowledgements

We thank the patients and their family members.

