## Supplementary figures and images for "*PHOX2B* polyalanine repeat mutation has a profound impact on the transcriptome of neuronal progenitor cells in Haddad syndrome"

### Suppl Fig. 1

**Suppl Figure 1:**

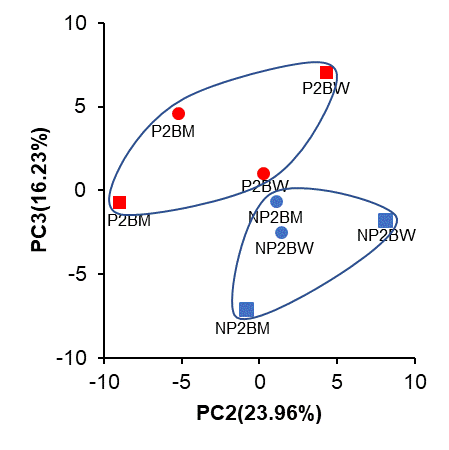

**Suppl Figure 1:** PCA depicting separate clustering of PHOX2B+ and PHOX2B-.
