## Supplementary material for "*PHOX2B* polyalanine repeat mutation has a profound impact on the transcriptome of neuronal progenitor cells in Haddad syndrome": Suppl. Fig 2

**Suppl Figure 2**

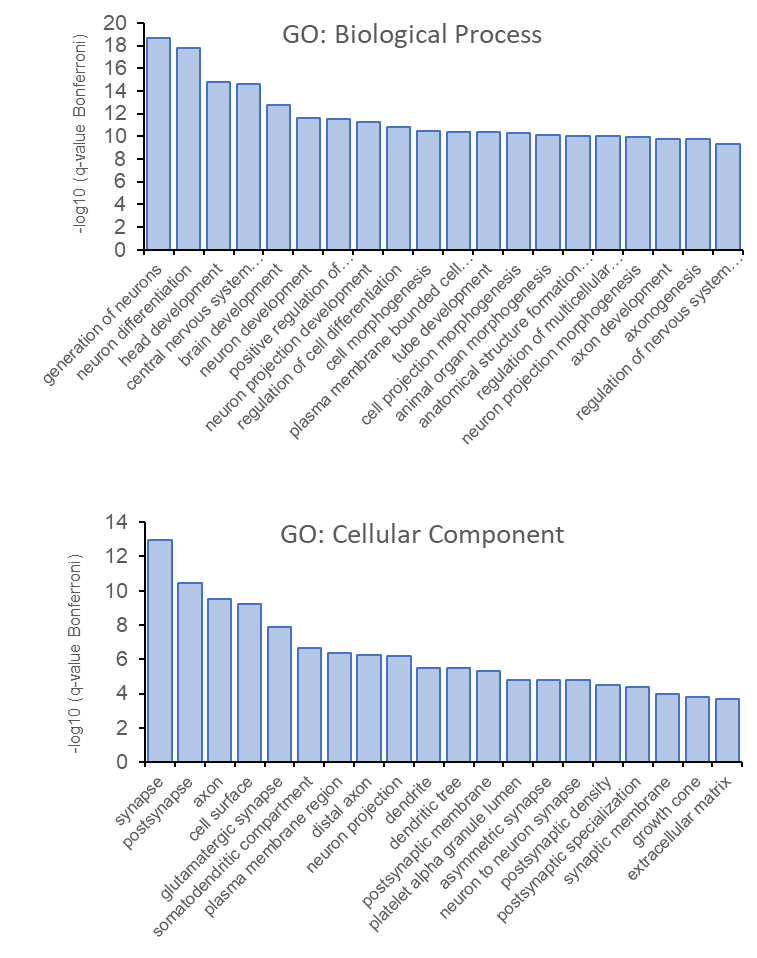

**Suppl Figure 2:** A) GO-biological process and (B) GO-Cellular component of the top DEGs between NP2BW vs P2BW.
